# PanACoTA: A modular tool for massive microbial comparative genomics

**DOI:** 10.1101/2020.09.11.293472

**Authors:** Amandine Perrin, Eduardo P.C. Rocha

## Abstract

The study of the gene repertoires of microbial species, their pangenomes, has become a key topic of study in microbial evolution and genomics. Yet, the increasing number of genomes available complicates the establishment of the basic building blocks of comparative genomics. Here, we present PanACoTA (https://github.com/gem-pasteur/PanACoTA), a tool that allows to download all genomes of a species, build a database with those passing quality and redundancy controls, uniformly annotate, and then build their pangenome, several variants of core genomes, their alignments, and a rapid but accurate phylogenetic tree. While many programs building pangenomes have become available in the last few years, we have focused on a modular method, that tackles all the key steps of the process, from download to phylogenetic inference. While all steps are integrated, they can also be run separately and multiple times to allow rapid and extensive exploration of the parameters of interest. The software is built in Python3 and includes features to facilitate its future development. We believe PanACoTa is an interesting addition to the current set of comparative genomics tools, since it will accelerate and standardize the more routine parts of the work, allowing microbial genomicists to more quickly tackle their specific questions.

## 1 Introduction

Low cost of sequencing and the availability of hundreds of thousands of genomes have made comparative genomics a basic toolkit of many microbiologists, geneticists, and evolutionary biologists. Many bacterial species of interest have now over 100 genomes publicly available in the GenBank RefSeq reference database, and a few have more than ten thousand. This trend will increase with the ever decreasing costs of sequencing, the availability of long-read technologies, and the use of whole-genome sequencing in the clinic for diagnostics and epidemiology. As a result, researchers that would like to use available data are faced with extremely large amounts of data to analyze. Comparative genomics has spurred important contributions to the understanding of the organization and evolution of bacterial genomes in the last two decades [1] [2]. It has become a standard tool for epidemiological studies, where the analysis of the genes common to a set of strains - the core or persistent genome - provides unrivalled precision in tracing the expansion of clones of interest [3] [4]. The use of routine sequencing in the clinic will further require rapid and reliable analysis tools to query thousands, and soon possibly millions of genomes from a single species [5]. Population genetics also benefits from this wealth of data because one can now track in detail the origin and fate of mutations of genetic acquisitions to understand what they reveal of adaptive or mutational processes [6]. Finally, genome-wide association studies have been recently adapted to bacterial genetics, to account for variants in single nucleotide polymorphism and gene repertoires [7]. They hold the promise of helping biologists to identify the genetic basis of phenotypes of interest. Given the high genetic linkage in bacterial genomes, these studies may require extremely large datasets to detect small effects. More specifically, reverse vaccinology is also a noteworthy application of these pangenomics methods, to identify novel potential antigens among core surface-exposed proteins of a given clade [8].

The availability of large genomic datasets puts a heavy burden on researchers, especially those that lack extensive training in bioinformatics, because their analysis implicates the use of automatic processes, efficient tools, extensive standardization, and quality control. Many tools have been recently developed to make rapid searches for sequence similarity with excellent recall rates for highly similar sequences [9] [10] [11].

Other tools also provide methods to rapidly cluster large numbers of sequences in families of sequence similarity, to get the families common to a set of genomes, to align them, or to produce their phylogeny, four cornerstones of comparative genomics. A number of recent programs have recently been published that include some of these tools to compute bacterial pangenomes (for a review, see [12]). Many of these programs compute alignments and clusters of families using programs that are very fast. Some use tools that are known to sacrifice accuracy for very high speed, such as DIAMOND [9], USEARCH [13] and CD-HIT [14]. The latter is used, among others by Roary [15], which is currently the most popular tool to compute pangenomes, and Panaroo [16], a very recent tool aiming at reducing the impact of erroneous automated annotation of prokaryotic genomes. BPGA [17], using USEARCH or CD-HIT to cluster proteins, also provides some downstream analyses. PanX [18], which has an outstanding graphical interface, uses DIAMOND to search for similarities among genes.

More recently, SonicParanoid introduced the use of the highly efficient and accurate program mmseqs2 to build pangenomes, and PPanGGOLiN used the same tool to provide a method to statistically class pangenome families in terms of their frequency [19] [20] [21]. Some recent programs also use graph-based approaches to further refine the pangenomes, such as PPanGGOLiN and Panaroo [16]. For that matter, the analysis of a dataset of 319 *Klebsiella pneumoniae* genomes by both tools provide very similar results [16]. Some tools, such as PIRATE [22] have also been recently developped to cluster orthologues between distant genomes. However, all these programs lack some or all of initial and final steps that are essential in comparative genomics, including download, quality control, alignment and phylogenetic inference. This spurred the development of PanACoTA (PANgenome with Annotations, COre identification, Tree and corresponding Alignments). To take advantage of the vast amount of genomic information publicly available, one needs six major blocks of operations. (1) Gather a set of genomes of a clade automatically. This requires some quality control, to avoid drafts with an excessive number of contigs. It is also often convenient to check that the genomes are not too redundant, to minimize computational cost and biases due to pseudo-replication. On the other side, it is important to check that genomes are neither too unrelated, to eliminate genomes that were misclassed in terms of bacterial species (or the taxonomic organisation of relevance). (2) Define *a priori* an uniform nomenclature and annotation, without which the calculation of pangenomes and core genomes becomes unreliable for large datasets. (3) Produce the pangenome, a matrix with the patterns of presence absence of each gene family in the set of genomes, using an accurate, simple, and fast method. (4) Use the pangenome to identify sets of core or persistent genes. (5) Produce multiple alignments of the gene families of the core or persistent genomes. (6) Finally, produce quickly a reasonably accurate phylogeny of the set of core/persistent genes. These four collections of data, pangenome, core genome, alignments, and phylogenetic tree, are the basis of most microbial comparative genomics studies. At the end of this process, the researcher can produce more detailed analyses, specific to the questions of interest, which often lead to changes such as including/excluding taxa, changing the limits of sequence similarity, increasing alignment accuracy, or rebuilding phylogenies using different methods. Such re-definitions can be achieved more efficiently when pipelines are modular and allow to re-start the analyses at several key points in the process.

Considering the current availability of pipelines for microbial comparative genomics, we have built one that is modular, easy to setup, uses state-of-the-art tools, and allows simple re-use of intermediate results. The goal was to provide a pipeline that allows to download all genomes from a taxonomic group and make all basic comparative genomics work automatically. The pipeline is entirely built in a single language, Python v3, and uses modern methods to facilitate its future maintenance and to limit unwanted behaviour. PanACoTA is freely available under the open source GNU AGPL license. Here, we describe the method and illustrate it with an analysis of two datasets of 225 complete and 3980 complete or draft genomes of *Klebsiella pneumoniae*. This species is interesting for our purposes because there are many genomes available and it has a very open pangenome [23]. The first dataset describes a situation where sequence quality is usually high, and the second illustrates how the method scales-up to a very large dataset where sequence is of lower quality or genes are fragmented due to lack of complete assembly. The procedure is detailed in the Methods section, whereas the illustration of its use, and how it changes in relation to key options in the two datasets, is detailed in the Results section.

## 2 Methods

PanACoTa is implemented in 6 sequential modules, described in the six sections below. It was designed to allow the use of a module without requiring the use of previous one(s). This allows to start or stop at any step and re-run an analysis with other parameters (see Figure 1).

**Figure 1:**
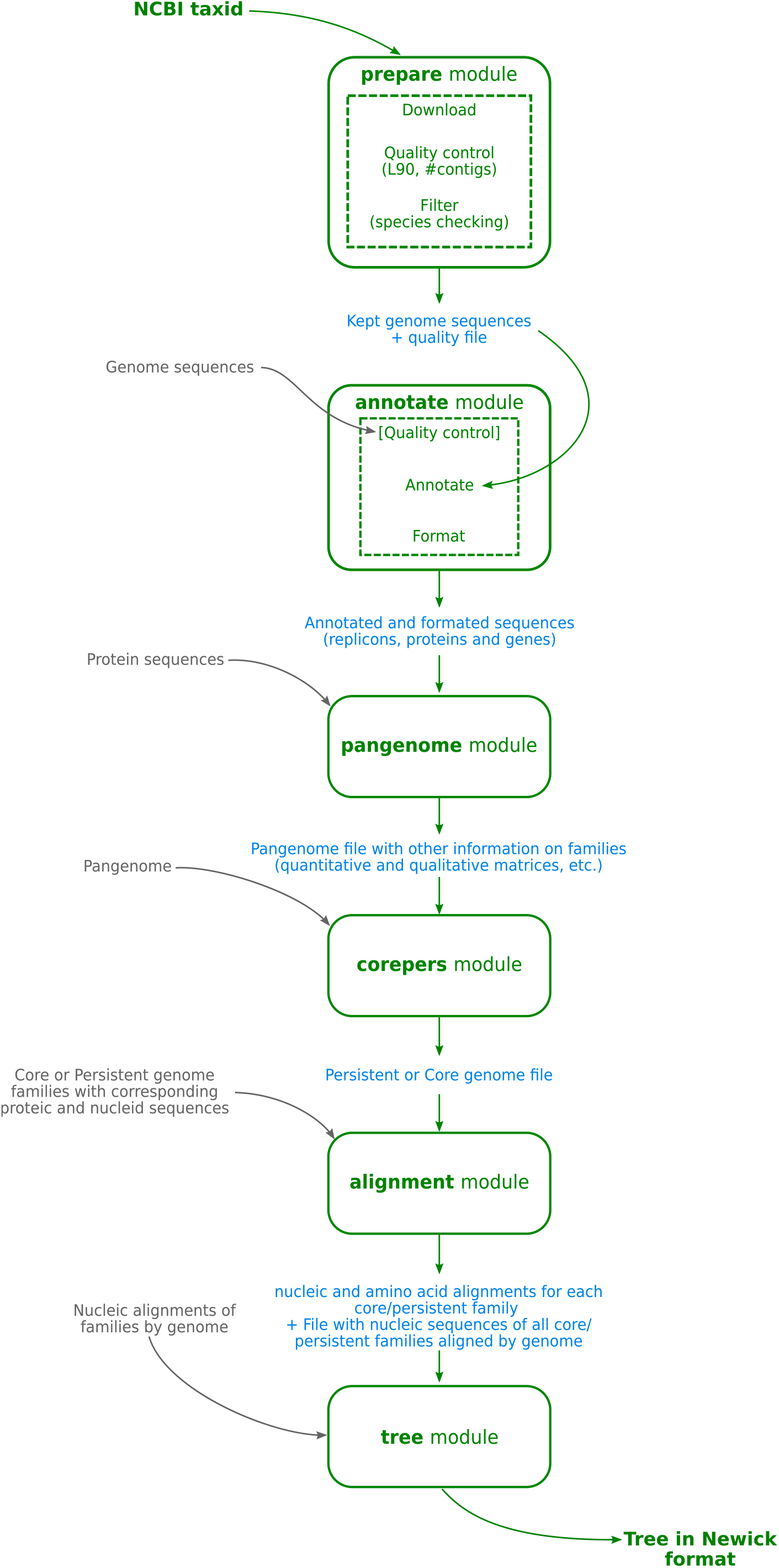
Overview of PanACoTA method

### 2.1 Datasets

The first module of PanACoTA - `prepare` - allows to fetch all genomes from a given NCBI taxonomy ID. This uses scripts from `ncbi_genome_download` library (https://github.com/kblin/ncbi-genome-download). PanACoTA retrieves the corresponding compressed non-annotated fasta files of the genomes from the NCBI ftp.

In this paper, we use two datasets of *Klebsiella pneumoniae* genomes to illustrate how PanACoTA functions. The first one, called hereafter DTS1, contains all complete and draft genomes downloaded from the NCBI refseq database on October 2018 10^th^. The second dataset, DTS2, is the subset of DTS1 containing only the complete genomes (genomes with `assembly_level = Complete Genome`, based on the NCBI summary file). This initial database was then given to a quality control procedure, described in the next section. At the end, this module ouputs a database with the genomes admissible after this control step: 3980 genomes for DTS1, with a subset of 225 complete genomes for DTS2. The accession numbers of all the genomes are indicated in Table S1.

### 2.2 Quality control procedure

The goal of this module is to remove genomes that do not conform with two types of basic requirements in terms of assembly and taxonomy. It is done by the `prepare` module after downloading the genomes, or by the `annotate` module before the annotation step (if the user did not use the `prepare` module). This latter option is useful when the goal is to analyse a predefined list of genomes, some of which are eventually not available in GenBank (e.g. in-house sequencing). The module receives as input a set of fasta sequences.

The first goal of this control procedure is to filter genomes in terms of sequence quality. Since there is usually no standard description of the quality of the sequence assembly in RefSeq genomes, the program infers it from the sequences. First, it is common usage to put stretches of ‘N’ to separate contigs in a same fasta sequence. To have a better idea of the sequence quality, and be able to do the analysis more efficiently, we first split sequences at each stretch of at least 5 ‘N’ (this number of 5 can be changed by the user) to get one fasta entry per contig. Assuming that the user is analyzing genomes from the same species, those genomes should have relatively similar characteristics in terms of number of contigs and length. Hence, PanACoTA first calculates two key measures for each genome: the total number of contigs, and the L90 (the minimum number of contigs necessary to get at least 90% of the whole genome). Very high values of these two variables are usually an indication of low quality of sequencing or assembling. They often result in the annotation of numerous truncated genes that spuriously increase the size of pangenomes (because a real gene is split into numerous open reading frames that are classed in distinct families). These poorly assembled genomes also complicate studies of comparative genomics whenever studying genetic linkage is important, because they are dominated by small contigs. The thresholds can be specified by the user. Values by default are set to less than 1000 contigs and L90 lower than 100. Genomes exceeding one of these values are excluded from the rest of the analysis.

The second part of the procedure is a filter dedicated to remove redundant and miss-classified genomes. This is done based on the genetic distance between pairs of genomes, as calculated by Mash [24]. We chose Mash for this distance filtering step because it can be computed very fast and is accurate for closely related genomes. Mash reduces each genome sequence to a sketch of representative k-mers, using the MinHash technique [25]. It then compares those sketches, instead of the full sequences. This output Mash distance `D` strongly correlates with alignment-based measures such as the Average Nucleotide Identity (ANI) which is based on whole-genome sequence comparisons using the blast algorithm [26]: *D* ≈ 1 – *ANI*. For ANI in the range of 90–100%, the correlation with Mash distance is even stronger when increasing the sketch size. Since pangenomes are typically computed for a single bacterial species, we are here using Mash to discriminate genomes having at least 94% identity. A few recent programs have been published showing slightly more accuracy than Mash, but we found them too slow for the use as a systematic filter. For example, using 15 cores, FastANI [27] requires around 1h15 to compare all pairs of 200 genomes (40,000 pairwise comparisons), where Mash with a sketch size of 10^6^ does the task in less than 3 minutes. Hence, a user requiring a finer grade study of ANI may wish to post-analyse the data from FastANI, instead of running the `prepare` module. However, it is impractical to make all pairwise analyses of very large datasets where one often needs to perform millions of pairwise comparisons.

Bacterial species are usually defined as groups of genomes at more than 94% identity [28], and this will typically be used as the upper value for D (`max_mash_dist` is a modifiable parameter that is fixed by default at 0.06). On the other extreme, genomes with very high similarity (corresponding to low Mash distances) provide very similar information. They can be excluded to lower the computational resources required for the analysis and to diminish eventual oversampling of certain clades, which could lead to biased results. PanACoTA sets `min_mash_dist` to 10^-4^ by default, but this parameter can also be specified by the user. This distance represents one point change every ten genes on average and may be close to the sequencing and assembling accuracy of many draft genomes.

The two procedures, quality control and Mash filtering, are linked together. The information on the number of contigs and L90 is useful to chose the genome that is kept between a pair of very similar genomes. In summary, the control procedure works as follows:

- Genomes with an excessively high number of contigs or L90 are excluded.
- Genomes are primarily sorted by increasing L90 value, and secondarily by increasing number of contigs to produce a list ordered in terms of quality.
- The genomes are compared with Mash. For that, the first genome of the ordered list (the one with best quality) is compared to all the others. The ones which do not obey to the distance thresholds are discarded. The procedure then passes to the subsequent genome in the ordered list (if not rejected before), compares it to all remaining genomes, and discards those not respecting the thresholds. The process continues until the ordered list is exhausted.

The output of this module is a database with the genomes that passed the two steps of the quality control procedure. PanACoTA also provides a file listing the discarded genomes and why they were discarded.

### 2.3 Annotation

The annotation of the genomes is done by the `annotate` module of PanACoTA. The input is a database of fasta sequences, from the `prepare` module or directly provided by the user. If no information is given on the quality control of those genomes (number of contigs and L90), this quality control is done here (see previous section for more information on the quality control step).

The goal of this module is to provide a uniform annotation of the gene positions (and functions) across the dataset. PanACoTA annotates all genomes with Prokka [29]. The latter uses Prodigal [30] to identify gene positions. It then adds functional annotations using a series of programs, including BLAST+ [31] to search for homologs in a database of proteins taken from Uniprot and HMMER3 [32] to search for proteins hitting selected profiles from TIGRFAM [33] and PFAM [34]. All annotated sequences are renamed using a standard sequence header format. The header of each gene contains 20 characters and provides human readable information on the genome and contig of the gene, its relative position in the genome, and if it is at the border of a contig (see Figure 2).

**Figure 2:**
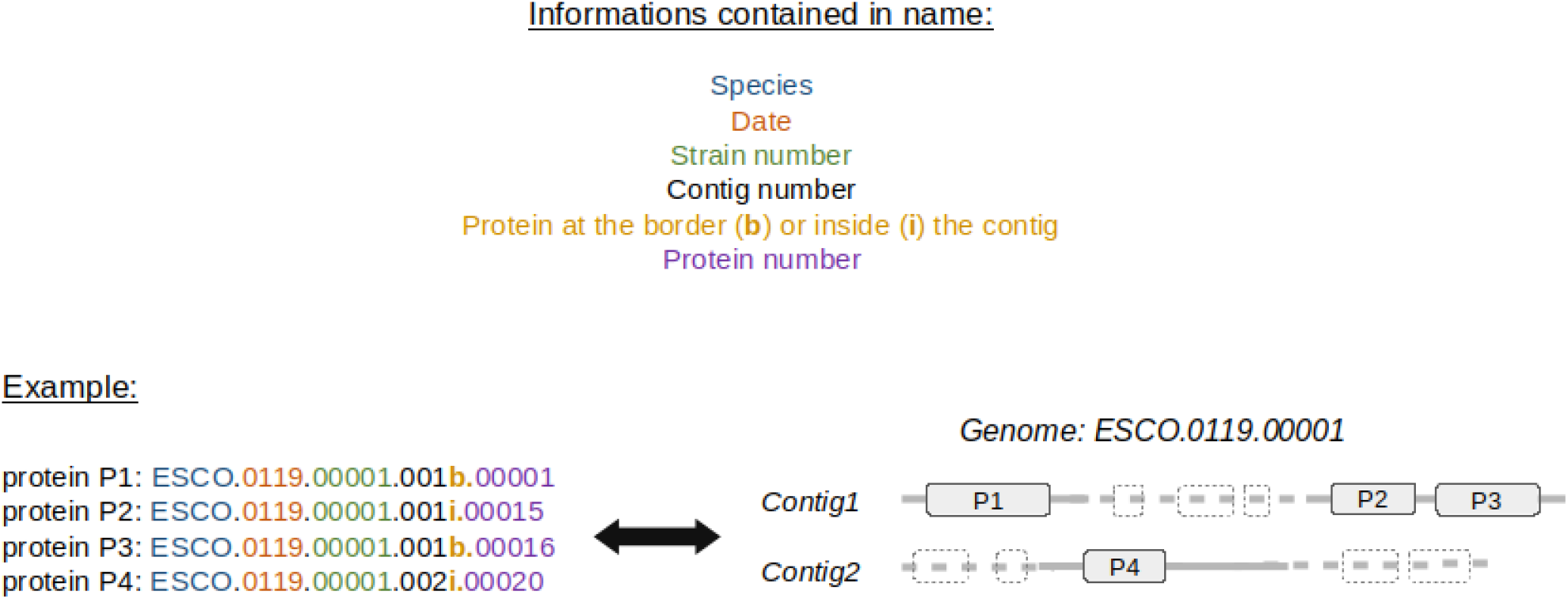
Description of the standard output header format for proteins annotated by PanA-CoTA.

If the user does not need the functional annotation, the module gives the possibility of running only the gene finding part, i.e. only running Prodigal. For very large datasets it is much faster to use this option and annotate a posteriori only one gene per family of the pangenome using Prokka or more complete annotation systems like InterProScan [35]. The output of this step consists in five files per genome: the original sequence, the genes, the proteins (all in fasta format), a gff file containing all annotations and a summary information file.

### 2.4 Identification of the pangenome

The pangenome is computed with the `pangenome` module of PanACoTA. The input is the set of all proteins from all genomes, e.g. the sequence files of the ‘Proteins’ folder generated by the `annotate` module.

The pangenome is the set of all protein families in the genomes. Its calculation involves comparisons between all pairs of proteins, i.e. its complexity is to the square of the number of genes (and thus of genomes). To generate a reliable pangenome in a reasonable time, PanACoTA calls the MMseqs2 suite [20]. The `mmseqs search` module has a very good speed/sensitivity trade-off. In order to reduce time, it uses 3 consecutive search stages, with increasing sensitivity and decreasing speed. Everything is highly parallelized and optimized on multiple levels. The first step filters up to 99.9% of the sequences by eliminating high dissimilarities, i.e. sequences not having at least two consecutive kmer matches. The second step filters out another 99% of the remaining sequences using an ungapped alignment. This leaves a small amount of sequences to process with an optimized version of the Smith-Waterman alignment, where only scores are calculated, and not the full alignments.

We used the `mmseqs cluster` module included in MMseqs2 suite, with the default `Cascaded clustering` option. This module works in two main steps. It first clusters proteins using linclust [36], a linear time protein sequence clustering algorithm as a prefilter. Then, the representative sequences of this first step are handled by the `mmseqs search` module, and clustered according to its result. This second step is repeated three times, each time with a higher sensitivity at the `mmseqs search` algorithm module.

For the clustering stage, PanACoTA uses the `Connected component` mode, because it has provided results consistent with our previous methods. This mode uses transitive connections to merge pairs of homologous genes: all vertices accessible via a BFS algorithm are members of a cluster. Let’s define the graph made from all pairwise comparisons between proteins as follows: each node is a protein and there is an edge between 2 proteins if they are similar (similarity beyond the given threshold). Then, two proteins are in the same family if we can find a path from one to the other in the graph. If desired, the user can choose any of two other clustering modes (`Greedy Set cover`, or `Greedy incremental`) using a dedicated parameter while launching the PanACoTA pangenome module. Importantly, the tuning of the options of mmseqs2 allows the sequence similarity analyses to be exceedingly fast or extremely sensitive [20]. In PanACoTA the user can change the key parameters `-min-seq-id` and `-cluster-mode`, and re-run the `mmseqs cluster` module to explore their effect on the results. More specific mmseqs2 parameters have, for the time being, to be used with the standalone version of the program.

The output of this step is a pangenome file containing one line per family, with the list of all its members. PanACoTA also provides the quantitative and qualitative matrices of the pangenome, as well as a tabular file giving an overview of each family composition.

Note that, here, we do not take into account synteny between genes in the genomes, as we think that, for draft genomes, this has a limited interest. By exploring the families generated, we found very few and non-significant differences. However, if the user has very well assembled genomes, or has any particular reason to account for synteny, some tools have been developped, like panOCT [37] [38], SynerClust [39] or PANINI [40].

If the user wants to do genome-wide association studies, the output qualitative matrix can be directly used as input for TreeWAS [41].

### 2.5 Identification of core and persistent genomes

The classification of gene families present in a large number of taxa is done by the corepers module of PanACoTA. The input is a pangenome file, like the one generated by the pangenome module. In early studies, the pangenome matrix was used to identify the gene families present in all genomes in a single copy: the core genome. However, the increase of the number of genomes in the dataset tends to decrease drastically the size of the core genome. This is because sequencing or annotation errors as well as rare deleterious polymorphism in the populations lead to the rapid decrease of the number of core genes with the increase in the number of input genomes. To overcome this problem, one now commonly identifies the persistent genome. A family is in the persistent genome if it contains members from at least N% of the genomes. N is defined by the user. The default value is 95% for datasets with more than 1000 genomes. The persistent genome is more robust to rare (true or artifactual) variants. On the other hand, if the goal of computing the persistent genome is to make a phylogenetic tree or analyze population genetics data, one may wish to produce sets with different thresholds. Indeed, a too high value will lead to a small set of persistent genes with few gaps that may be enough to infer a robust phylogeny. On the other side, this will exclude many gene families from other types of analyses, like detection of positive selection or recombination events. The definition of persistent genome may also vary, depending on the subsequent use of the data. PanACoTA defines three types of persistent genomes (see Figure 3):

- Strict-persistent: a family that contains exactly 1 member in at least `N`% genomes (`N` = 100 means it is a core-family). This definition is particularly practical to reconstruct phylogenies without having to handle the existence of multiple copies per genome.
- Mixed-persistent: a family where at least `N`% of the genomes have exactly 1 member, and other genomes have either 0, either several members in the family. This definition is intermediate between the other two, i.e. it includes the strict-persistent and is included by the multi-persistent.
- Multi-persistent: a family with at least one member in `N`% of the genomes. This definition is interesting to analyse patterns of diversification of nearly ubiquitous protein families.

**Figure 3:**
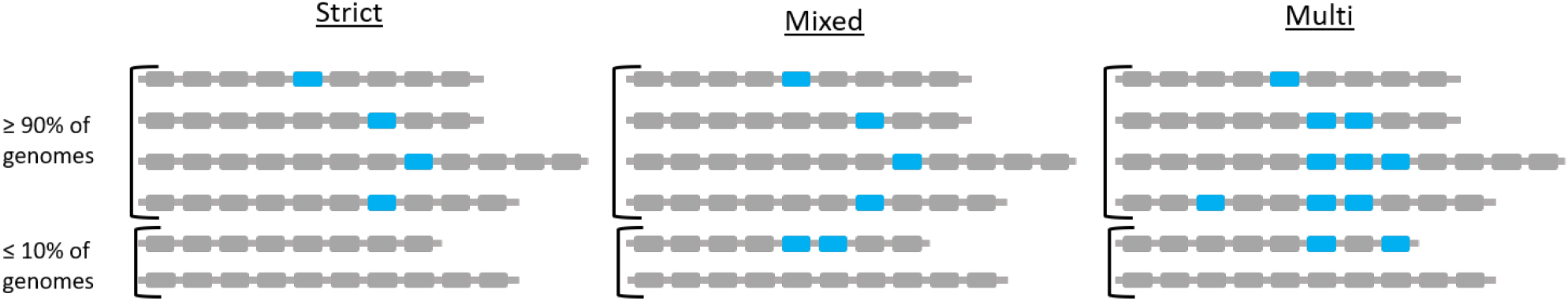
Different types of persistent genomes proposed by PanACoTA, with a threashold of N = 90%.

The mixed and multi-persistent definitions are useful to include the gene families with variable numbers of copies in a small (mixed) or large (multi) number of genomes when studying the evolution of nearly ubiquitous gene families. It can be useful when one protein was split in several parts in a few genomes because of sequencing or assembly error(s). This protein family is discarded from the strict-persistent genome, while included in the mixed (and multi) persistent genomes.

One should note that the module `corepers` does the re-analysis of the pangenome and therefore it does not use a reference genome whose choice can be questionable. Re-running the module is very fast, because it only requires the re-analysis of the pangenome matrix. Hence, it is easy and fast to re-run the module with different parameters in different analysis to check how they change the final result.

The output of this module is a file containing the persistent families of proteins.

Note that if the user wants to identify the persistent genome using a statistical approach rather than using fixed thresholds, the gff file generated by `annotate` module is compatible with PPanGGOLiN [21]. This software generates a persistent genome corresponding to our multi-persistent version of the persistent genome (multigenic families are allowed).

### 2.6 Multiple alignments of the persistent gene families

The alignment of the persistent gene families is done by the `align` module of PanACoTA. Its input is a persistent genome coming from the `corepers` module, or independently provided by the user. When using the strict-persistent genome, all genes are aligned. When using the other definitions of persistent genomes, some genomes can lack a gene or have it in multiple copies. To produce phylogenetic trees from these alignments, such cases must be handled beforehand. When a genome lacks a member or has more than one member (mixed or multi persistent) of a given gene family, PanACoTA adds a stretch of gaps (‘-’) of the same length as the other aligned genes. Adding a few “-” has little impact on phylogeny reconstruction. For example, it has been showed that adding up to 60% of missing data in the alignment matrix could still result in informative alignments [42]. In our experience, when this approach is applied to within-species pangenomes, it usually incorporates less than 1% of gaps. The effect of missing data should thus be negligible relative to the advantage of using the phylogenetic signal from many more genes (i.e. in contrast to using the strict-persistent genome). Alignments are more accurate when done at the level of the protein sequence. This has the additional advantage of producing codon-based nucleotide alignments that can be used to study selection pressure on coding sequences. Hence, PanACoTA translates sequences, aligns the corresponding proteins and then back-translates them to DNA to get a nucleotide alignment. This last step constitutes in the replacement of each amino acid by the original codon. Hence, at the end of the process, the aligned sequences are identical to the original sequences.

PanACoTA does multiple sequence alignment using MAFFT [10] as it is often benchmarked as one of the most accurate multiple alignment programs available and one of the fastest [43]). It has options that allow to make much faster alignments, at the cost of some accuracy, to handle very large datasets. This loss of accuracy is usually low for very similar sequences as it is the case of orthologous gene families within species, and means that PanACoTA can very rapidly align the persistent genome.

This module returns several output files: the concatenate of the alignments of all families to be used for tree inference, and, for each core/persistent genome family, a file with its gene and protein sequences aligned.

### 2.7 Tree reconstruction

The phylogenetic inference is done with the `tree` module of PanACoTA. It uses as input the alignments of the `align` module or any other alignments in Fasta format.

This is the part that takes most time in the entire pipeline, because the time required for phylogenetic inference grows very fast with the size of the dataset. Even efficient implementations of the maximum likelihood analyses scale with the product of the number of sites and the number of taxa, which is a problem in the case of large datasets (thousands of taxa, with more than ten thousands sites for each one). PanACoTA proposes several different methods to obtain a phylogeny: IQ-TREE [44], FastTreeME [45], fastME [46] and Quicktree [47]. Whatever the software used, the `tree` module takes as input a nucleotide alignment in Fasta format (like, for example, the output of `align` module), and returns a tree in Newick format. According to its needs, the user can choose one of these methods to infer its phylogenetic tree. These trees can be used to build more rigorous phylogenetic inference using methods that are more demanding in computational resources, e.g. by changing the options of IQ-TREE.

### 2.8 Implementation and availability

`PanACoTA` was developed in Python3, trying to follow the best practices for scientific software development [48] [49]. For that, the software is versioned using git, allowing the tracking of all changes in source code during PanACoTA’s development. It is freely distributed under the open-source AGPL licence (making it usable by many organizations) and can be downloaded from https://github.com/gem-pasteur/PanACoTA.

Hosting it on GitHub allows for issue tracking, i.e. users can report bugs, make suggestions or, for developers, participate to the software improvement. To provide a maintainable and reliable software, we set up continuous integration process: each time a modification is pushed, there is an automatic software installation checking, unit tests are done, and, if necessary, an updated version of the documentation is generated, as well as an update of the singularity image on Singularity Hub.

As introduced just before, we also provide a complete documentation, including a step by step tutorial, based on provided genome examples, so that the user can quickly get started. It also contains more detailed sections on each module, aiming at helping users to tune all parameters, in order to adapt the run to more specific needs. This documentation also includes a ‘developer’ section, addressed to developers wanting to participate in the project.

During its execution, PanACoTA provides logging information, so that user can see realtime execution progress (a `quiet` parameter is also proposed for users needing empty stdout and stderr). This also provides log file(s) to keep track on what was ran (command-line used, time stamp, parameters used etc.).

## 3 Results and discussion

All execution times mentioned in this section correspond to wall clock time on 8 cores (except when the number of cores is given). A summary of all execution times can be found in Table 1.

**Table 1:** Summary of execution times by (sub)module

| MODULE | STEP | DTS1 (3980 genomes) | DTS2 (225 genomes) |
| --- | --- | --- | --- |
| prepare | downloading | 1h (5805 genomes) | 3min (266 genomes) |
|  | quality control | <4min | ~15sec |
|  | filter | 20min | ~1min |
| annotate | with Prokka | 5 days | 10h |
|  | with Prodigal | 6h | 30min |
| pangenome |  | 30min | 1min |
| corepers (1 core) |  | 1min | 5sec |
| align | strict persistent | 3h | 10min |
|  | mixed-persistent | 7h | 11min |
| Tree (IQ-TREE)<br>(28 cores) | strict-persistent | 7h (40GB RAM) | 3min10 |
|  | mixed-persistent | 24h (90GB RAM) | 3min30 |

### 3.1 Download and preparation of genome sequences

The first module of PanACoTA was used to download all genomes of *Klebsiella pneumoniae* using the TaxID 573. It took approximately 1h to download the 5805 *Klebsiella pneumoniae* genome sequences (including 266 complete genomes). We used the module `annotate` to make the quality control of those 5805 strains. For this study, we used as thresholds: L90<100 and number of contigs<999. The computation of L90, number of contigs and genome size, and the subsequent procedure of discarding genomes according to the thresholds took less than 4 minutes. This step discarded 233 draft genomes, leaving 5572 for further analysis (see Figure 4). When the threshold on the number of contigs was decreased by half (number of contigs<500), only 52 more genomes were removed (see Figure 4b). To define the best thresholds to the analysis, the user can preview its dataset quality with a ‘dry-run’ of the `annotate` module. Then, the user can launch the real analysis, from `prepare` or `annotate` with the adapted thresholds.

**Figure 4:**
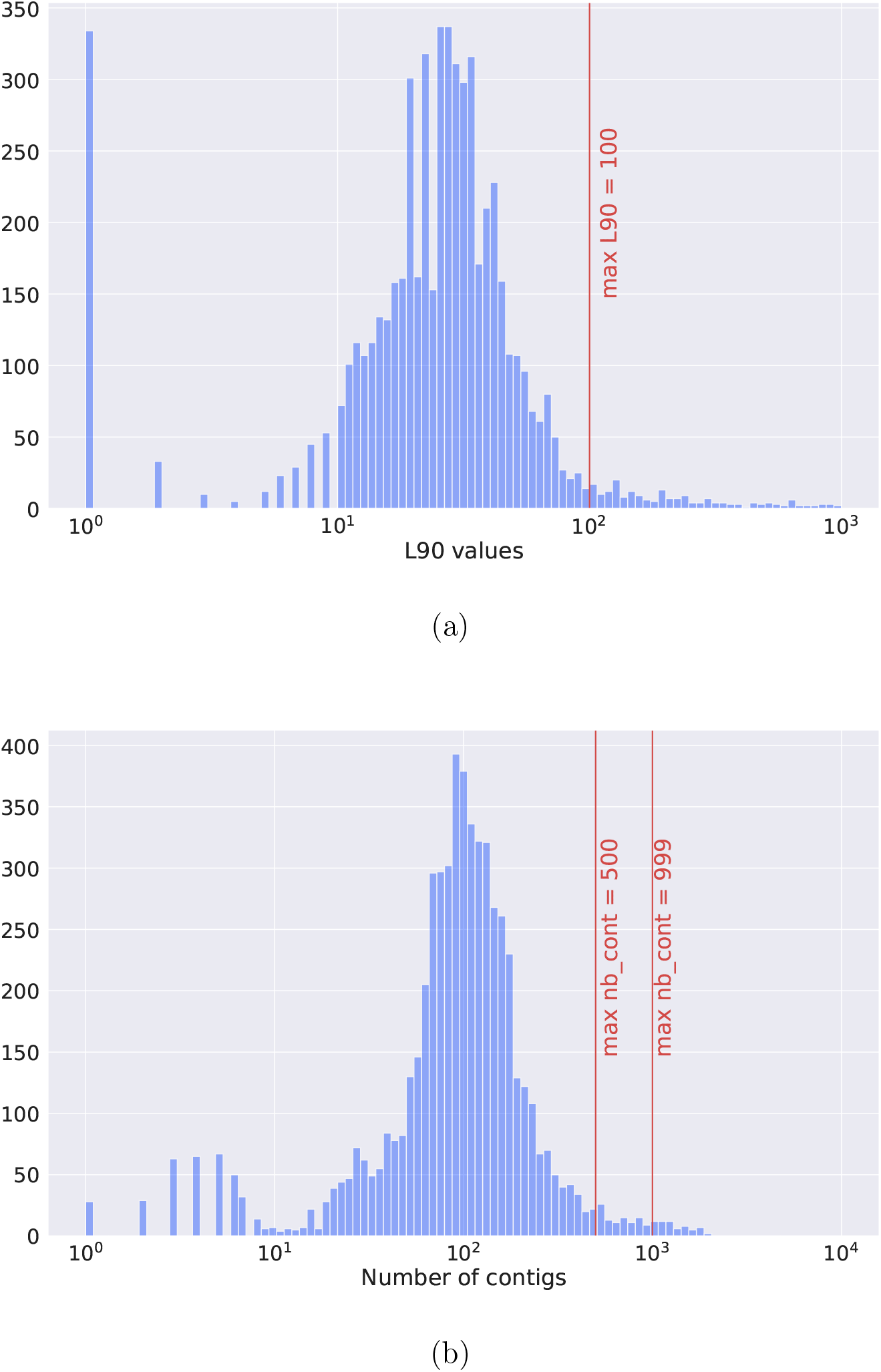
Histograms describing the features of the 5805 *K. pneumoniae* genomes downloaded from Refseq. (a) Distribution of L90 values. (b) Distribution of the number of contigs per genome.

The analysis of genetic distances across pairs of genomes is performed by Mash (K-mer size of 21 (default), and sketches of at most 10000 non-redundant min-hashed k-mers). A total of 1592 genomes (including 41 complete genomes) did not respect the distance thresholds (`max_mash_dist` = 0.06 and `min_mash_dist` 1e^-4^)). The vast majority (1448) were too similar to other genomes, whereas 144 were too distant from other strains to be regarded as bona fide *Klebsiella pneumoniae* genomes (figure 5).

**Figure 5:**
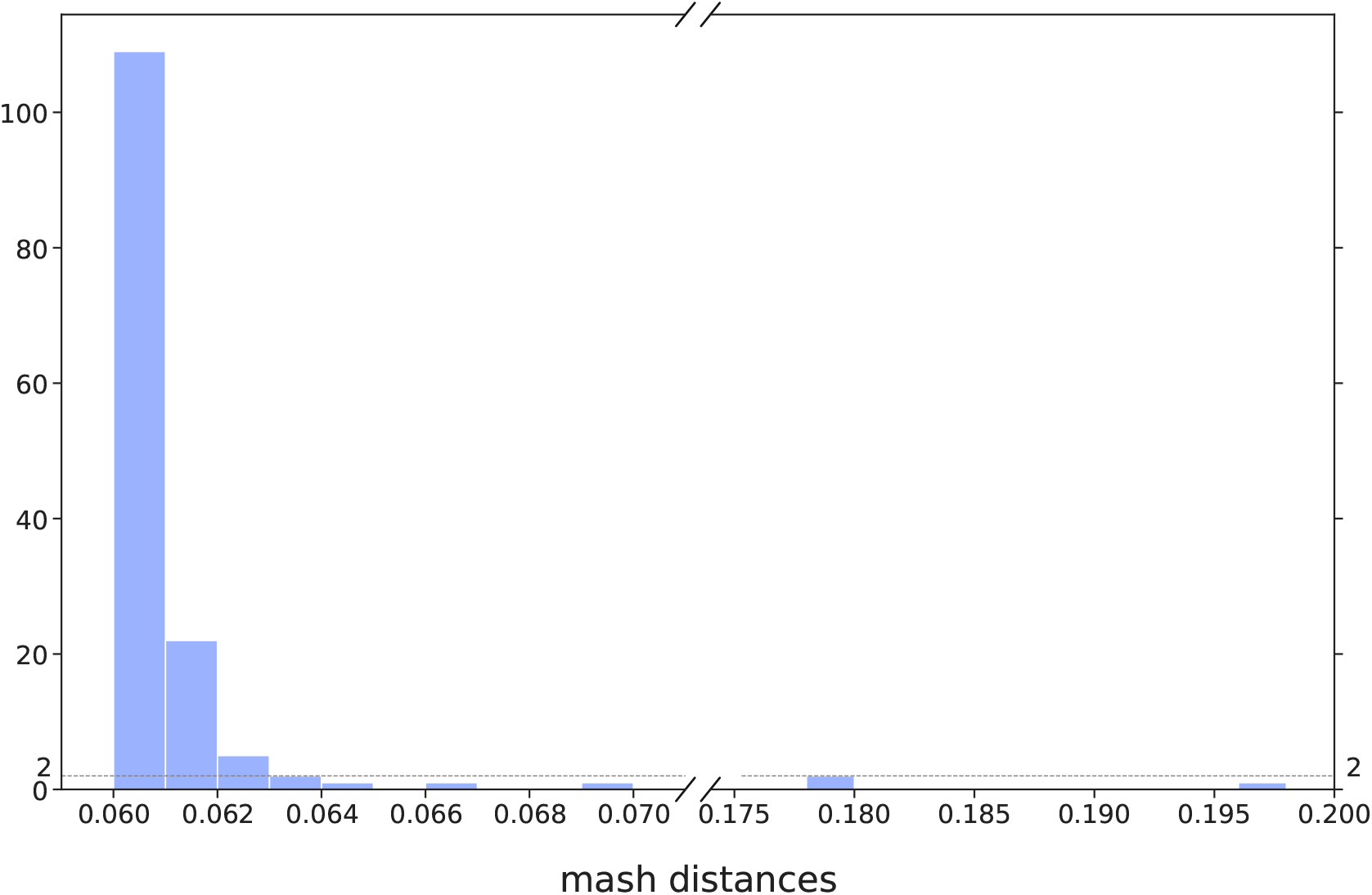
Distribution of Mash distances for the 5572 genomes respecting the L90 and number of contigs thresholds, but having a Mash distance higher than the threshold (0.06).

Some species can even be defined with narrower ANI values. For example, to identify bona fide *Klebsiella pneumoniae* genomes, Kleborate (https://github.com/katholt/Kleborate) uses Mash to compare the given assembly to a curated set of *Klebsiella* assemblies from NCBI. It considers a Mash distance of ≤ 0.01 as a strong species match, and a Mash distance between 0.01 and 0.03 as a weak match. With our DTS1, Kleborate would have only removed 22 more genomes, that it identifies as *Klebsiella quasipneumoniae subspecies similipneumoniae*. Our method, which is designed for any species, is thus quite consistant with Kleborate results regarding the specific case of *K. pneumoniae* genomes.

Three genomes showed an ANI less than 84% identity, meaning they may not even be from the same genus, which emphasizes the necessity of this kind of analysis before computing a pangenome. They were removed from the analysis (GCF_900451665.1, GCF_900493335.1 and GCF_900493505.1). Finally, these filters left 3980 genomes in the analysis, with an average of 5307 genes per genome, which will be called the reference database DTS1. Among them, there are 225 complete genomes that form the dataset DTS2 (see Figure 6).

**Figure 6:**
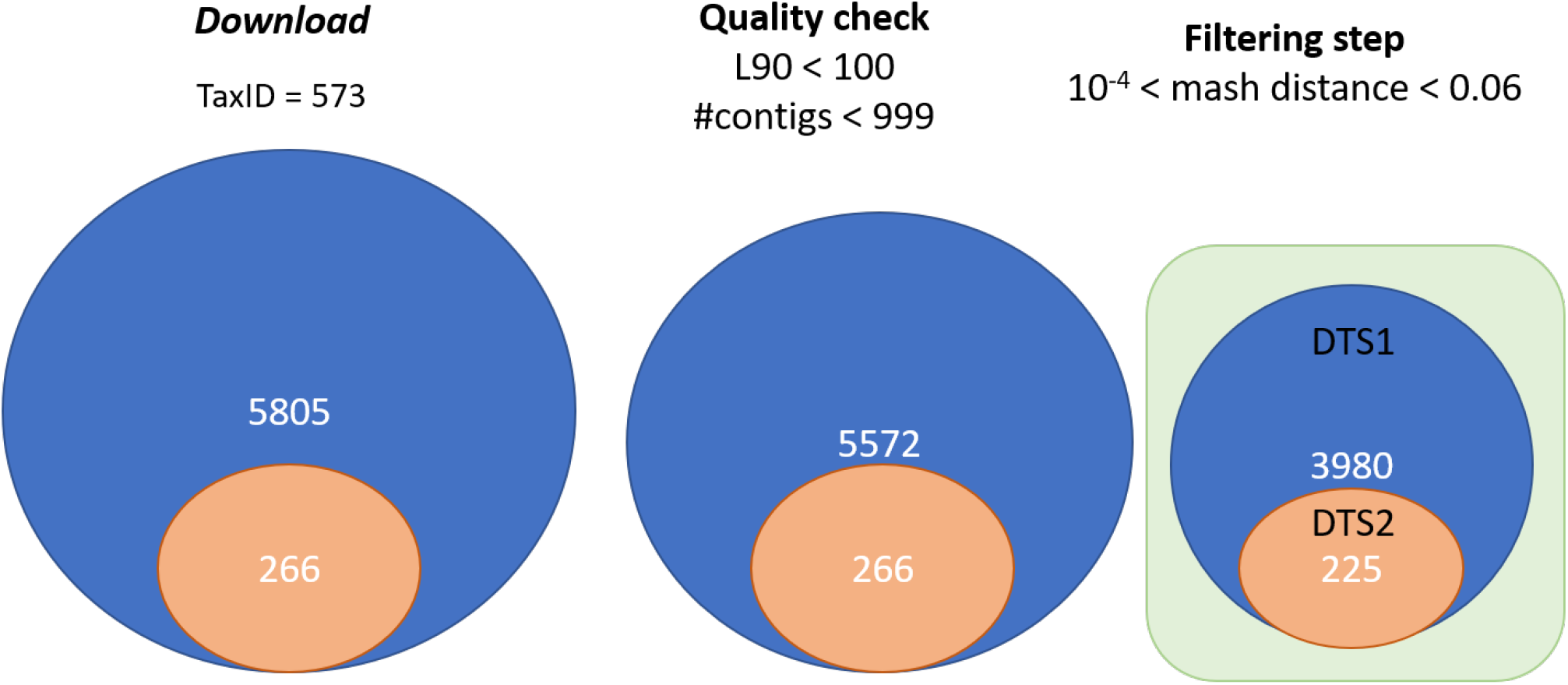
Summary of the procedure to construct DTS1 and DTS2.

The functional annotation part is by far the slowest of the first modules. On a typical 5 Mb genome, giving 2 cores to Prokka, gene finding takes around 40 seconds and functional annotation around 7 minutes.

The annotation of the genomes with Prokka 1.11 took approximately 1min 50s per genome, i.e. around 5 days for the whole dataset. For comparison, we re-did the analysis with the option of restricting the analysis to prodigal 2.60. This analysis took less than 6h in total (annotation + formatting of all 3980 genomes), which corresponds to an average of 6 seconds per genome.

In general, this shows that making a simple syntactic annotation leads to a considerable gain of time. Assuming that genes from the same pangenome family have similar functions, one can annotate one protein per family at the end of the process and save considerable time.

### 3.2 Building pangenomes

We used MMseqs2 Release 11-e1a1c. The 3980 DTS1 genomes contain 20,765,062 proteins. It took less than 30min to create the protein database in the MMseqs2 format, cluster them (with at least 80% identity and 80% coverage of query and target, and other parameters kept as default), and retrieve the pangenome matrices. The pangenome of the smaller DTS2 dataset of 225 genomes, 1,190,485 proteins, was computed in less than one minute. The DTS1 pangenome has 86607 families. Among them, 35348 (40%) are singletons (found in a single genome), which is concordant with values observed in *Escherichia coli* [50]. In DTS2 we found 24473 families, including 8975 (37%) singletons.

The comparison of these two pangenomes is interesting because it reveals the robustness of the method to changes in sampling size, as summarized in Figure 7. A total of 2147 families contain only members present in both DTS1 and DTS2. This means that, for those families, even with all proteins of DTS1, only proteins from DTS2 were clustered together. Among those families, 2122 are exactly the same in both pangenomes, whereas only 25 families were split in the DTS1 pangenome family relative to the DTS2 pangenome. In that case, they are split, most of the time, into two different families of DTS1. This shows that the clustering procedure is quite robust to the addition of a very large number of genomes.

**Figure 7:**
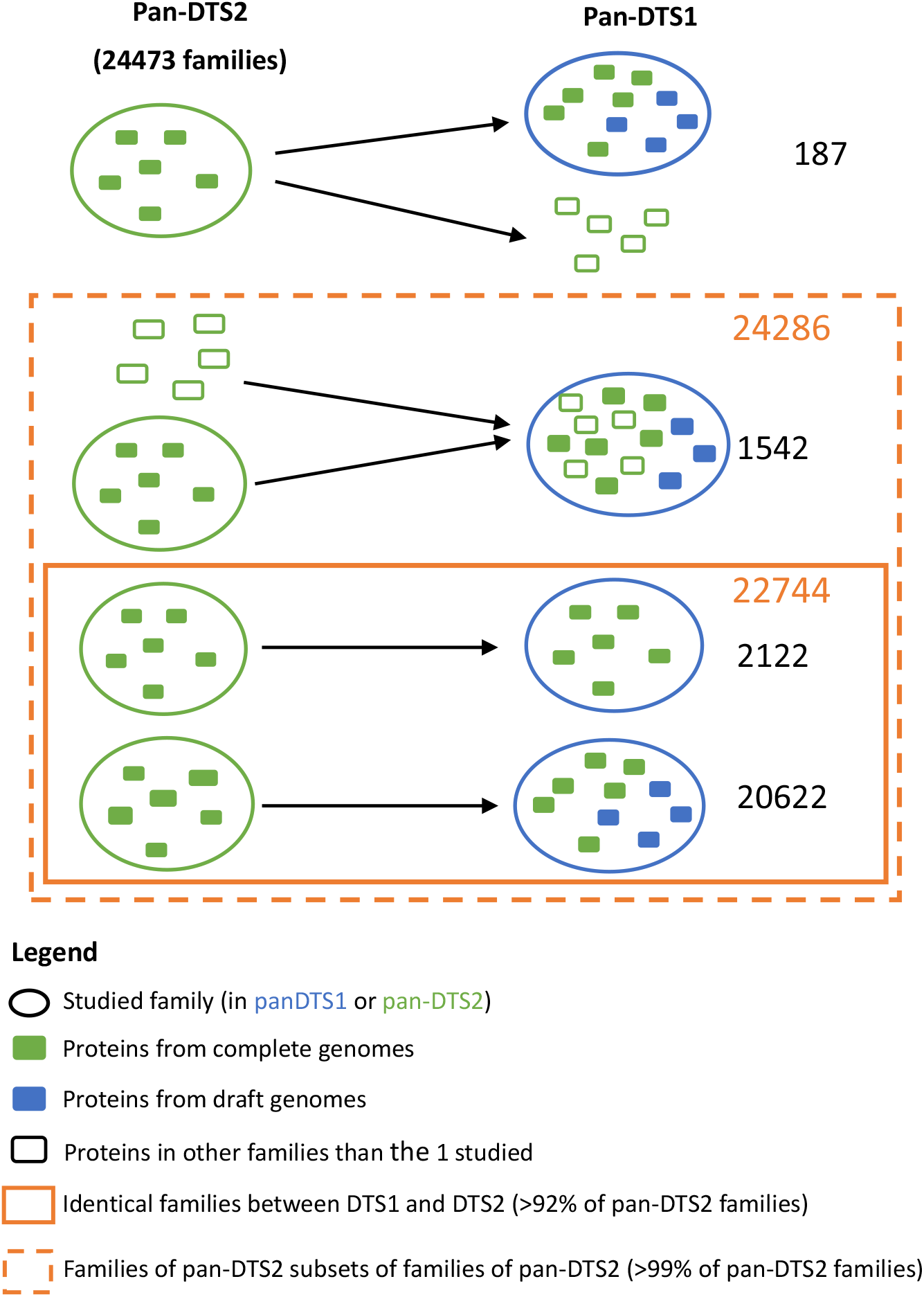
Comparison of the pangenomes generated by PanACoTA for both DTS1 and DTS2.

Most important, we observed a total of 22744 families (that is more than 92% of all DTS2 families) that are identical between the independent analysis of the DTS1 and DTS2 pangenomes. Identical here means that the DTS2 pangenome gene family is included in a DTS1 pangenome gene family, and the other members of this DTS1 pangenome family are only members of genomes not present in DTS2. Furthermore, around half of the remaining families from the DTS2 pangenome are included in a DTS1 pangenome gene family, which contains a few other proteins from DTS2 genomes. Finally, only 187 gene families of the DTS2 pangenome were split into 2 or 3 different families of DTS1 pangenome. In other words, 24286 families (more than 99%) of DTS2 pangenome are subsets of DTS1 gene families. In conclusion, the construction of pangenome families is robust to large variations in the number of input genomes (see Figure 7).

### 3.3 Core and persistent genomes

This part of the analysis is very fast. Using only 1 core, it took around one minute to generate a core or persistent genome from DTS1 pangenome. PanACoTA provides a core genome and three different measures of persistent genome (see Figure 3). The strict-persistent genome corresponds to cases when the family is present in a single copy in 99% genomes and absent from the others. In DTS2, the set of complete genomes, the difference between the core and strict-persistent genome is appreciable (2238 versus 3295 families), i.e. the persistent genome is 50% larger (see Figure 8). The difference becomes huge when the analysis is done on the much larger and less accurate DTS1 dataset, where the two datasets vary by more than one order of magnitude (79 versus 1418 families). In such large datasets of draft genomes the analysis of the core genome is not very useful.

**Figure 8:**
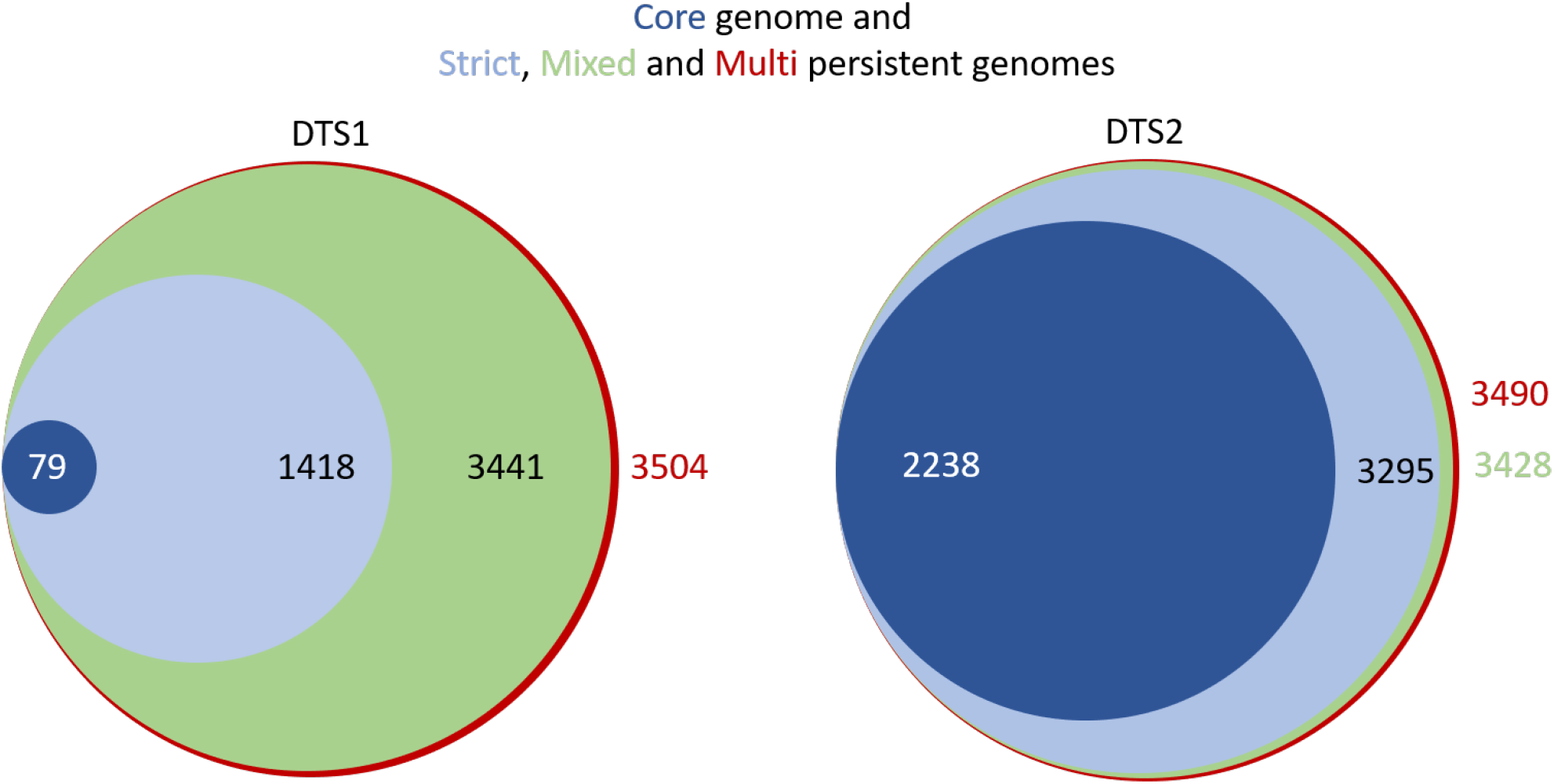
Comparison of the sizes of the core genome and the 3 different types of persistent genomes, for both DTS1 and DTS2. Areas of circles are proportional to the size of the dataset.

The mixed-persistent genome includes the families present in a single copy in 99% genomes and present (potentially in several copies) or absent from the others. It includes the strict-persistent genome and is not much larger than the latter in the small DTS2 dataset. Yet, the difference is much larger in the DTS1 dataset (see Figure 8). While the mixed-persistent genome is 65% percent of the average genome in DTS1, the strict-persistent is only 27% percent in the same dataset. This shows the relevance of using definitions of the core genome adapted to the dataset in order to build robust phylogenetic trees or to analyse patterns of genetic diversification and natural selection.

Finally, PanACoTA also computes a multi-persistent genome that includes all gene families present in at least 99% genomes, independently of the copy number. It includes all the other sets and is not much larger than the mixed-persistent genome (see Figure 8). Yet, it includes interesting families. An analysis of these reveals many genes encoding regulators, transporters and enzymes that are nearly ubiquitous, but often present in multiple copies. As a rule, this definition is interesting to study gene families present in most genomes, but present in very different copy number. On the other hand, it is typically not very useful for phylogenetic inference. Since all these sets can be computed very rapidly, it’s straightforward to compute them all and use them for different types of analyses.

### 3.4 Phylogenetic tree inference

PanACoTA ran mafft v.7.467 using `--auto` option to align all families. For DTS1, it selected the FFT-NS-2 method, while for DTS2, it selected FFT-NS-i method. This was done with both the strict-persistent (1418 families) and the mixed-persistent (3441 families). It took 3h (resp. 7h) to align all the families of the strict-persistent (resp. mixed-persistent), giving, for each one, the input file for tree inference.

For tree inference, PanACoTA used IQ-TREE multicore version 2.0.6, with `-fast` option. For the tree based on the alignment of the strict-persistent (3980 sequences, 1418 families corresponding to 1,438,179 positions), it took around 7h on 28 cores, requiring 38GB of RAM. For the tree based on the alignment of the mixed-persistent (3980 sequences, 3441 families corresponding to 3,393,006 positions), it took 24h using 28 cores, requiring 88GB of RAM.

We then wished to understand the differences in phylogenetic inference in terms of the method used to define the persistent genome (strict and mixed persistent). We computed the patristic distance matrix for each tree and a Pearson correlation test showed that they are strongly correlated (cor = 0.99138, p < 2.2e-16). This shows that the distances provided by the two methods are very similar. Hence, if the strict persistent is large enough to generate a phyogenetic tree, it provides adequate distances between genomes. Aligning all mixed persistent families would just take much more time, for a similar result. However, if one is interested in having a robust tree topology, one should use the larger (and computationally costlier) dataset. Indeed, the analyses of Robinson-Foulds distance with R `phangorn` package shows a branch-weighted distance of 0.43 and an absolute distance of 2892 [51]. This is because some lineages of *K. pneumoniae* account for a large fraction of the data and these parts of the tree require long informative multiple alignments to produce accurate topologies. Accordingly, the differences in topology between the trees using the DTS2 dataset, which have much larger average branch lengths, show much smaller values of topological distances between the two datasets of persistent genome (RF=78, wRF=0.027).

Some researchers use methods to detect recombination in genomes, remove the recombination tracts, and then redo the analyses. This can be done outside PanACoTA by querying the multiple alignments before proceeding to the phylogenetic inference. Yet, previous results have shown that such procedures tend to distort phylogenetic inference at a larger extent than simply using all the information in the multiple alignments [52] [53]), and this explains why we have not included such an option on PanACoTA.

## 4 Conclusion

PanACoTA is a pipeline for those wanting to test hypotheses or explore genomic patterns using large scale comparative genomics. We hope that it will be particularly useful for those wishing to use a rapid, accurate and standardized procedure to obtain the basic building blocks of typical analyses of genetic variation at the species level. We built the pipeline having modularity in mind, so that users can produce multiple variants of the analyses at each stage. We also paid particularly care with the portability and evolvability of the software. These two characteristics, modularity and evolvability, will facilitate the implementation of novel procedures in the future.

## Data Availibility

The two datasets of *Klebsiella pneumoniae* genomes used to illustrate PanACoTA were downloaded from the NCBI refseq. Their accession numbers are indicated in Supplementary Table S1. PanACoTA source code is freely available from https://github.com/gem-pasteur/PanACoTA under AGPLv3 license. More information in 2.8.

## Acknowledgements

We thank Marie TOUCHON, Matthieu HAUDIQUET and Rémi DENISE for comments, suggestions, and bug reports, Yoann DUFRESNE for his help on optimizing some parts of the code, and Blaise LI for his help with Singularity. This work was partly supported by the ANR Salmo_Prophages (ANR-16-CE16-0029), the Inception program (PIA/ANR-16-CONV-0005), and the Equipe FRM (EQU201903007835).

## Notes

### Competing Interest Statement

The authors have declared no competing interest.

